# Polysome fractionation analysis reveals features important for human nonsense-mediated mRNA decay

**DOI:** 10.1101/2020.03.08.981811

**Authors:** James P. B. Lloyd, Courtney E. French, Steven E. Brenner

**Author notes:** ARC Centre of Excellence in Plant Energy Biology, University of Western Australia, Perth, Australia. Department of Paediatrics, School of Clinical Medicine, University of Cambridge, CB2 0QQ, UK.

## Abstract

Nonsense-mediated mRNA decay (NMD) is a translation-dependent mRNA surveillance pathway that eliminates transcripts with premature termination codons. Several studies have tried defined the features governing which transcripts are targeted to NMD. However, these approaches often rely on inhibiting core NMD factors, which often have roles in non-NMD processes within the cell. Based on reports that NMD-targeted transcripts are often bound by a single ribosome, we analyzed RNA-Seq data from a polysome fractionation experiment (TrlP-Seq) to characterize the features of NMD-targeted transcripts in human cells. This approach alleviates the need to inhibit the NMD pathway. We found that the exon junction complex (EJC) model, wherein an exon-exon junction located ≥50 nucleotides downstream of a stop codon is predicted to elicit NMD, was a powerful predictor of transcripts with high abundance in the monosome fraction (bound by a single ribosome). This was also true for the presence of an upstream open reading frame. In contrast, as 3’ UTR lengths increase, the proportion of transcripts that are most abundant in the monosome fraction does not increase. This suggests that either longer 3’ UTRs do not consistently act as potent triggers of NMD or that the degradation of these transcripts is mechanistically different to other NMD-targeted transcripts. Of the ribosome-associated transcripts annotated as “non-coding”, we find that a majority are bound by a single ribosome. Many of these transcripts increase in response to NMD inhibition, including the oncogenic SHNG15, suggesting many might be NMD targets. Finally, we found that retained intron transcripts without a premature termination codon are over-represented in the monosome fraction, suggesting an alternative mechanism is responsible for the low level of translation of these transcripts. In summary, our analysis finds that the EJC model is a powerful predictor of NMD-targeted transcripts, while the presence of a long 3’ UTR is not.

## Introduction

Gene expression is a complex set of interconnected processes including transcription, alternative splicing, RNA stability and translation. Nonsense-mediated mRNA decay (NMD) is a translation-dependent RNA decay mechanism that targets transcripts with an early stop codon (Peltz et al., 1993; Ishigaki et al., 2001). Several models have been proposed to explain what marks a stop codon as authentic, where termination of translation occurs normally, versus a premature termination codon (PTC), which leads to NMD in animals, yeast and plants (Nagy & Maquat, 1998; Amrani et al., 2004; Bühler et al., 2006; Behm-Ansmant et al., 2007; Kérenyi et al., 2008; Singh et al., 2008; Hogg & Goff, 2010; Drechsel et al., 2013; Lloyd et al., 2018). Alternative splicing has a well known role in altering the coding potential of transcripts, but can also alter the translation efficiency of the transcript (Floor & Doudna, 2016; Weatheritt et al., 2016; Blair et al., 2017; Fagg et al., 2017; Reixachs-Sole et al., 2019) or change the stability of transcripts by targeting them to NMD (Jumaa & Nielsen, 1997; Hilleren & Parker, 1999; Lejeune et al., 2001; Lewis et al., 2003; Lareau et al., 2007; Ni et al., 2007; Floor & Doudna, 2016; Reixachs-Sole et al., 2019).

The EJC model, also known as the 50nt rule, has been proposed to describe how a stop codon is recognized as a PTC in mammals, which leads to degradation by NMD. Reporter constructs expressed in mammalian cells have determined that exon-exon junctions located ≥50-55 nt downstream of a stop codon would trigger NMD (Nagy & Maquat, 1998; Zhang et al., 1998a; Zhang et al., 1998b). Such a stop codon was termed a Premature Termination Codon; we designate this as PTC_50nt_. Exon-exon junctions are associated with exon junction complexes (EJCs), which are deposited on a transcript during splicing, where the EJC marks the location of the spliced intron (Le Hir et al., 2000a; Le Hir et al., 2000b). Normally, the ribosome removes EJCs during the first round of translation (Gehring et al., 2009). However, if the EJC is associated with an exon-exon junction located ≥50-55 nt downstream of a stop codon, the ribosome does not remove the EJC and the EJC signals that the stop codon is premature (Le Hir et al., 2001; Lykke-Andersen et al., 2001; Kashima et al., 2006; Lopez-Perrote et al., 2016). Alternative models suggest that transcripts can be targeted to NMD due to the presence of a long 3’ UTR (Amrani et al., 2004; Bühler et al., 2006; Behm-Ansmant et al., 2007; Singh et al., 2008; Hogg & Goff, 2010; Bao et al., 2016; Fanourgakis et al., 2016; Ge et al., 2016; Kishor et al., 2019). Several intronless transcripts and transcripts with no intron in their 3’ UTR have been found to be targeted to NMD, and the long lengths of their 3’ UTRs have been reported to be responsible (Bühler et al., 2006; Kerényi et al., 2008; Yepiskoposyan et al., 2011; Toma et al., 2015; Balagopal & Beemon, 2017; Nyiko et al., 2017).

It has previously been reported that some NMD targets in *C. elegans* are mostly bound by a single ribosome (Barberan-Soler et al., 2009). By separating the cell’s RNA out on a mass gradient, fractions of RNA corresponding to transcript bound by a single ribosome (the monosome fraction) or transcripts bound by multiple ribosomes (the polysome fractions) can be isolated. In subsequent work, NMD targets are observed at greater levels in the monosome fraction than in the polysome fractions in yeast (Heyer & Moore, 2016). Additionally, predicted NMD-targeted transcripts were depleted from the polysome fraction, relative to the total cytoplasm or monosome fraction, in human cells (Sterne-Weiler et al., 2013; Floor & Doudna, 2016; Kim et al., 2017; Mencke et al., 2017). These findings are consistent with the evidence that many NMD targets are recognized in the pioneer round of translation (Ishigaki et al., 2001; Chiu et al., 2004). If the termination of translation triggers NMD, the first translating ribosome could trigger NMD (Ishigaki et al., 2001; Chiu et al., 2004; Barberan-Soler et al., 2009). It has been shown that the nuclear cap complex promotes NMD in mammalian cells (Hwang et al., 2010), further suggesting that NMD would occur before many ribosomes have loaded on the transcript, and often when only a single NMD-eliciting ribosome is bound. Therefore, transcripts that are most abundant in the monosome fraction are likely enriched for NMD targets.

Previous studies characterizing the features of transcripts that lead to NMD have often relied upon inhibition of NMD via direct inhibition of an NMD factor, or by inhibiting translation with drugs such as cycloheximide or emetine. All of these approaches will lead to changes in the transcriptome unrelated to NMD; for example, because NMD factors often have extensive roles in non-NMD processes (Azzalin & Lingner, 2006; Isken & Maquat, 2008; Lloyd, 2018).

In this study, we used the relative high abundance of NMD targets in the monosome fraction compared to the polysome fraction to test features previously linked to NMD: the EJC model, longer 3’ UTRs, and upstream open reading frames (uORFs). By characterizing the abundance of transcripts in different polysome fractions, we are able to avoid many of the secondary effects of long term NMD inhibition and find NMD targets in physiologically normal cells. We used an RNA-Seq dataset generated from polysome fractions (Transcript Isoforms in Polysomes-Seq; TrIP-Seq) by Floor and Doudna (2016). Floor and Doudna performed polysome fractionation of a human cell line (HEK293) on a sucrose gradient and then sequenced RNA in the monosome fraction, in polysome fractions representing two through seven ribosomes binding to a single transcript and in a polysome fraction representing eight or more ribosomes bound to a single transcript. We then tested which NMD linked features were associated with transcript isoforms most abundant in the monosome fraction. We found support for the EJC model acting as a potent trigger of NMD, as transcripts with an exon-exon junction ≥ 50nt downstream of a stop codon (designated 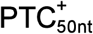) are much more relatively abundant in the monosome fraction than transcripts without a PTC_50nt_ (designated 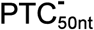). In contrast, we do not see a correlation between 3’ UTR length and relative abundance in the monosome fraction. In the monosome fraction, we also found enrichment of transcripts containing uORFs with a consensus Kozak sequence and transcripts annotated as non-coding RNAs, suggesting NMD could play an important role in the expression of these transcripts. Finally, we find that many retained intron transcripts are most abundant in the monosome fraction even when no premature termination codon is present. This suggests that a novel mechanism controls the sequestration of these transcripts in the monosome fraction, similar to what has been previously reported for retention in the nucleus of retained intron transcripts (Gohring et al., 2014; Boutz et al., 2015; Braun et al., 2017).

## Results

### Relatively high abundance of transcripts with a PTC_50nt_ in the monosome fraction supports the EJC model of NMD

NMD targets have previously been shown to be most abundant in the monosome fraction (Barberan-Soler et al., 2009; Sterne-Weiler et al., 2013; Floor & Doudna, 2016; Heyer & Moore, 2016). Motivated by this quality of NMD targets, we characterized transcript abundance across the polysome fractions for features associated with NMD. We re-analyzed polysome fractionated RNA-Seq data (Floor & Doudna, 2016) by quantifying a transcriptome assembled using transcript data after NMD inhibition, therefore capturing many more NMD targeted transcripts than in a reference transcriptome (French, 2016) (Supp Table 1). By the nature of this experiment, it is enriched for non-NMD targets because NMD targets are degraded and therefore low abundance. However, not introducing a global treatment to inhibit NMD allows us to examine potential NMD features without artefacts arising from these treatments. We examined in depth SRSF6 and hnRNP DL, which each have alternative splicing, leading to production of distinct transcript isoforms that are differentially sensitive to NMD (Lareau et al., 2007; Ni et al., 2007; French, 2016; Kemmerer et al., 2018). For the major SRSF6 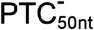 transcript isoform, 77% of the transcripts were bound by eight or more ribosomes and for the minor SRSF6 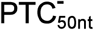 isoform 87% were bound by eight or more ribosomes (Fig 1A). This is in contrast to the 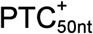 SRSF6 transcript isoform; only 6% of this transcript isoform was bound by eight or more ribosomes (Fig 1A). The majority of the SRSF6 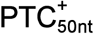 transcripts were bound by either a single ribosome (39%) or by two ribosomes (23%; Fig 1A). A similar pattern emerges for hnRNP DL, where 65% of one 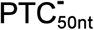 transcript isoforms was bound by eight or more ribosomes as was 36% of another, while 65% of the 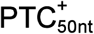 transcript isoform was bound by a single ribosome (Fig 1B). The relatively high abundance of 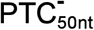 transcripts bound by eight or more ribosomes is consistent with these transcripts being productive and the substrate for most translation of these genes, in contrast, the relatively high abundance of 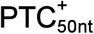 transcripts being bound by a single ribosome is consistent with these transcripts being NMD targets.

**Fig 1.**
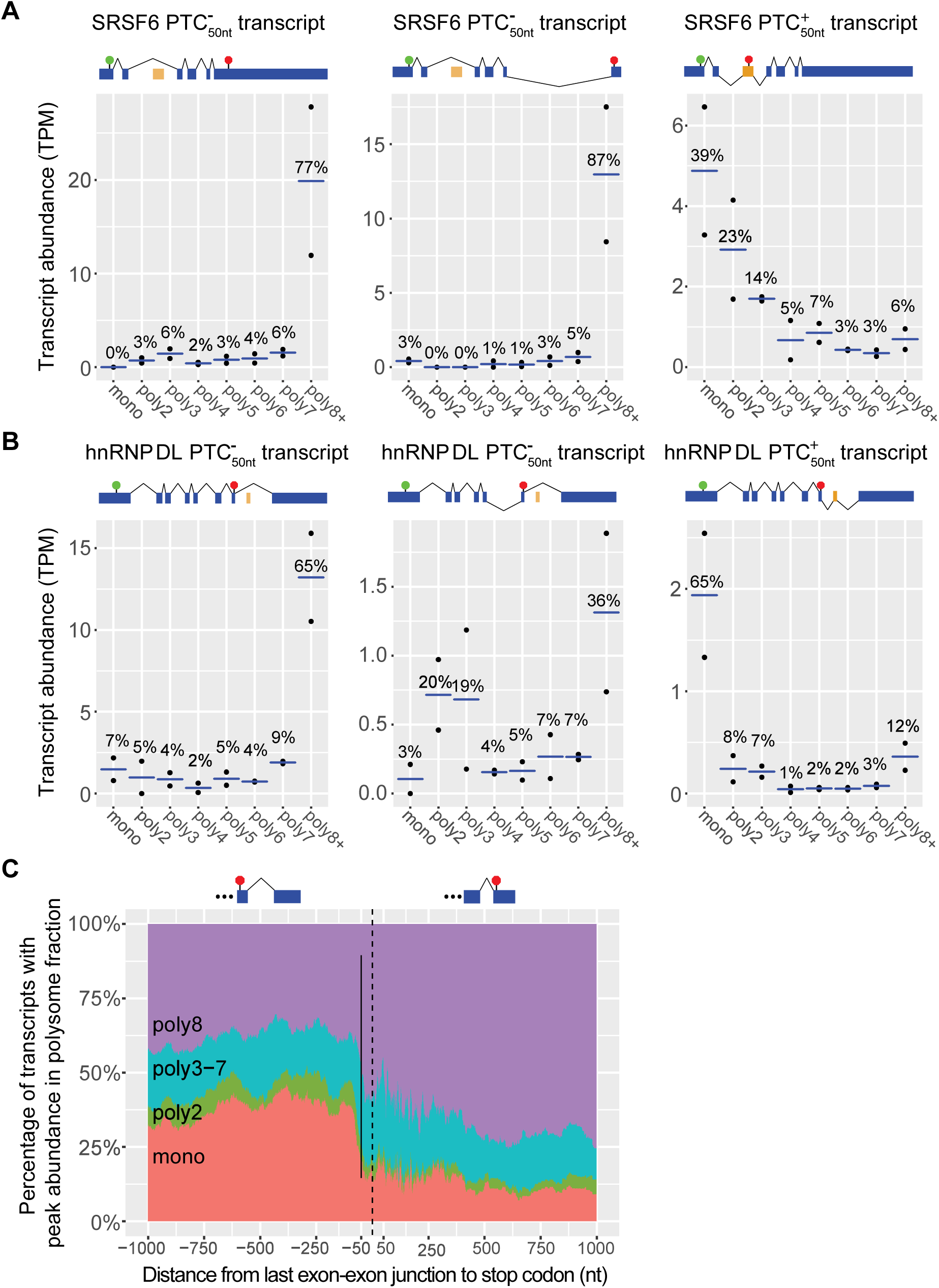
Transcripts with a PTC_50nt_ are enriched in the monosome fraction. (A) The abundance of SRSF6 transcripts in the different polysome fractions according to whether they have a PTC_50nt_. SRSF6 (ENSG00000124193) transcript IDs from left to right: 5_00146653 (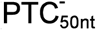 transcript), 5_00146654 (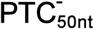 transcript), and 5_00146652 (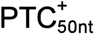 transcript). (B) The abundance of hnRNP DL transcripts in the different polysome fractions. hnRNP DL (ENSG00000152795) transcript IDs from left to right: 5_00037797 (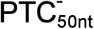 transcript), 5_00037799 (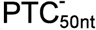 transcript), and 5_00037796 (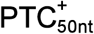 transcript). For (A) and (B), transcript abundance is measured in TPM normalized across the polysome fractions. Black dots represent the abundance in each of the two replicates and the blue bar represents the average abundance between the replicates. Blue exons are constitutively spliced exons. Orange exons are alternatively spliced exons. Circumflexes above the transcript model denote splicing of the major transcript isoform, while circumflexes below the transcript model represent alternative usage of a junction. (C) The distance between stop codon final exon-exon junction versus the proportion of transcripts most abundant in each polysome fraction. Abundance of transcripts found most predominately in monosome fraction increases sharply when the stop codon is more than 50 nt before the last exon-exon junction. Positive values on the right of the plot represent transcripts whose last exon-exon junction are located before the stop codon. Negative values located on the left of the plot represent transcripts whose last exon-exon junctions are located after the stop codon. Dotted line marks zero, where the first base of the stop codon is immediately after an exon-exon junction, while the solid bar marks −50 nt. Intronless transcripts are excluded from this plot. The proportion of transcripts most abundant in each polysome fraction at each position along the x axis is a average of a sliding window analysis, ensuring that each position represents at least 200 transcripts.

For each transcript isoform in the transcriptome, we measured the distance from the last exon-exon junction to the stop codon. We then examined how this related to peak abundance in the monosome fraction (Fig 1C). In agreement with the EJC model, we find a sharp increase in the proportion of transcripts mostly bound to a single ribosome when the last exon-exon junction is ≥50 nt downstream of the last exon-exon junction (Fig 1C). A relatively low proportion (15%) of 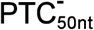 transcripts are most abundant in the monosome fraction (Fig 1C), while a relatively higher fraction (35%) 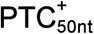 transcripts are most abundant in the monosome fraction (Fig 1C). 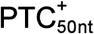 transcripts are also relatively enriched (7%) in the polysome 2 fraction relative to the 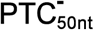 transcripts (3%) (Fig 1C). An example of this is the SRSF10 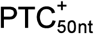 transcript (Supp Fig 1A). If the initiation rate is significantly high or the coding sequence significantly long, a second round of translation may start before the pioneering ribosome has terminated and initiated NMD; this would lead to two or more ribosomes binding to the NMD targeted transcripts and explain the enrichment of 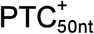 transcripts in the polysome 2 fraction (Fig 1C). Although per-gene initiation rate data are not available, we do know the sequence lengths of 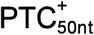 transcripts. By comparing the coding sequence length of 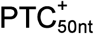 transcripts that are most abundant in the monosome fraction and the polysome 2 fraction (Supp Fig 1B), we find a small and non-significant difference in the lengths (average 213nt vs 258nt, Wilcoxon rank sum test p = 0.09). This suggests that longer coding sequences might partly account for why some NMD-targeted transcripts are most abundant in the polysome 2 fraction but other differences in these transcripts are the main determinants. Taken together, these data show that exon-exon junctions, when located ≥ 50 nt downstream of a stop codon are highly associated with transcript abundance in the monosome and polysome 2 fractions. The low number of ribosomes on transcripts with a PTC_50nt_ is consistent with the role of the EJC in targeting transcripts to NMD, consistent with the EJC model.

### Long 3’ UTRs are not correlated with relatively high abundance in the monosome fraction

If long 3’ UTRs generally act to target transcripts to NMD, we would expect to see a correlation between 3’ UTR length and proportion of isoforms whose highest abundance is in the monosome fraction. To explore this, we restricted our analysis to 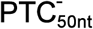 transcripts—since a PTC_50nt_ is already known to act as a potent trigger of NMD (Nagy & Maquat, 1998; Zhang et al., 1998a; Zhang et al., 1998b; Drechsel et al., 2013; Hurt et al., 2013; Kurosaki & Maquat, 2013; French, 2016; Lindeboom et al., 2016; Colombo et al., 2017; Lloyd et al., 2018; Hoeket al., 2019), and would confound the results. We found that increasing 3’ UTR length does not correlate with an increase in transcripts that have peak abundance in the monosome fraction (Fig 2A). Indeed, we see a gentle decline in the proportion of transcripts that have peak abundant in the monosome fraction as 3’ UTR length increases (Fig 2A). We also examined the profiles of transcripts whose 3’ UTRs have been reported to trigger NMD in reporter construct assays (Yepiskoposyan et al., 2011; Toma et al., 2015). We find that SMG5 has a flat abundance profile across the different polysome fractions (Fig 2B) and that SMG7 and MFN2 are most abundant in the polysome 8+ fraction (Fig 2B), which is indicative of highly translated transcripts. Taken together, these data show that transcripts with long 3’ UTRs are not enriched for peak abundance in the monosome fraction. For those targeted to NMD, this suggests differences in NMD of these transcripts relative to those with a PTC_50nt_.

**Fig 2.**
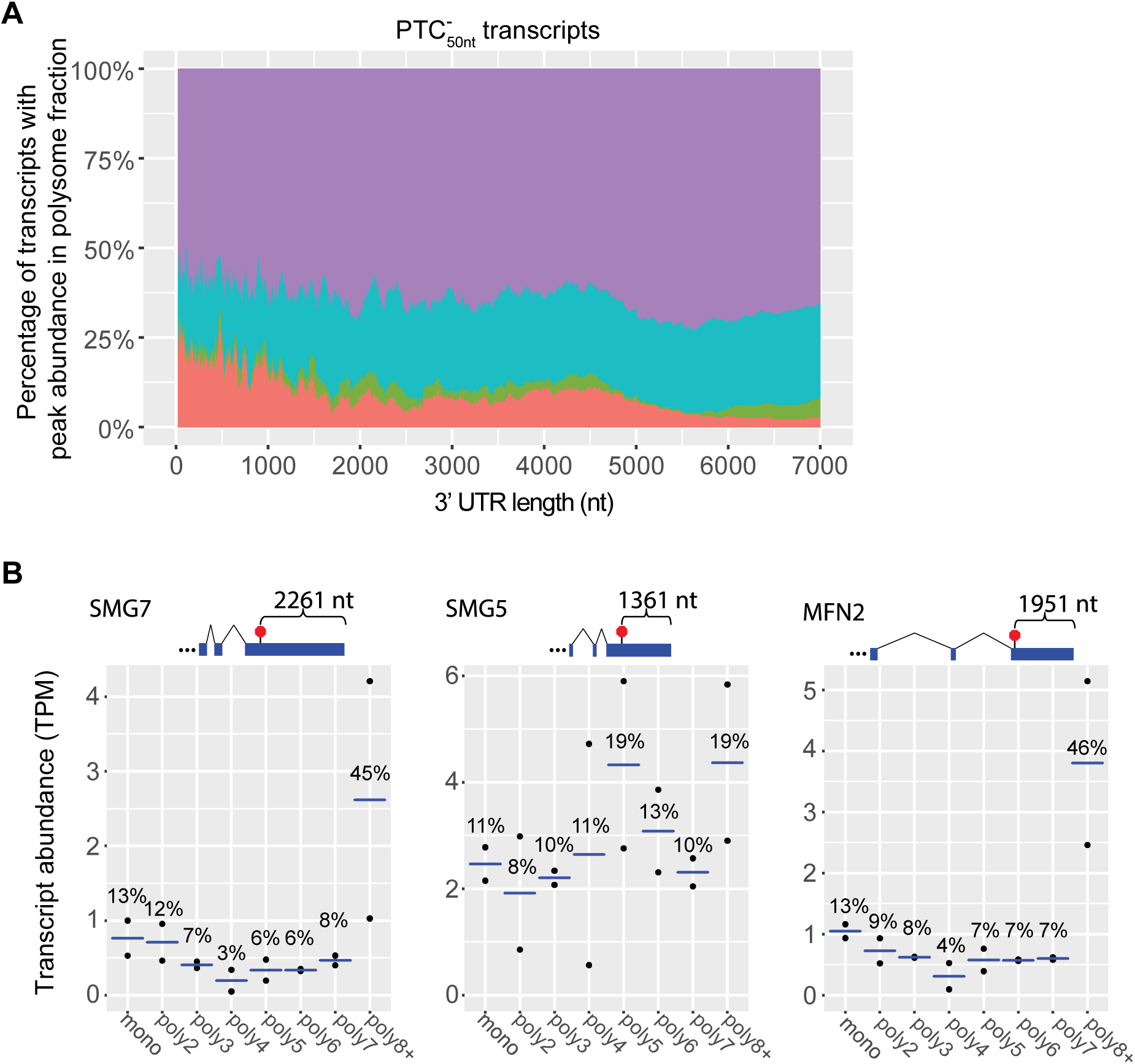
Transcripts with a long 3’ UTR are not enriched in the monosome fraction. (A) Length of transcript 3’ UTRs versus the proportion of transcripts most abundant in the different polysome fractions. The proportion of transcripts most abundant in different polysome fractions at each position along the x axis is a moving average by sliding window analysis, ensuring that each position represents at least 200 transcripts. (B) Example transcripts whose 3’ UTRs have been shown to target transcripts to NMD in reporter assays (graphical conventions follow Fig 1). SMG7 (ENSG00000116698) transcript ID: 5_00003622, SMG5 (ENSG00000198952) transcript ID: 5_00007602, and MFN2 (ENSG00000116688) transcript ID: 5_00000356.

### Physiologically relevant NMD target GADD45A is not most abundant in the monosome fraction

NMD is essential for normal development in many animals and removal of a core NMD factor leads to embryonic lethality in flies and mammals (Medghalchi et al., 2001; Metzstein & Krasnow, 2006; Weischenfeldt et al., 2008). The cause of the lethality in some animals has been linked to NMD targeting GADD45, which encodes a regulator of programmed cell death (Nelson et al., 2016). It is unclear what features of GADD45 target it to NMD, as it lacks a PTC_50nt_ and a long 3’ UTR but appears to be a direct NMD target (Nelson et al., 2016). In agreement with previous reports (Viegas et al., 2007; Huang et al., 2011; Tani et al., 2012; Nelson et al., 2016; da Costa et al., 2019), we found that GADD45A did increase after NMD inhibition (Fig 3A; Supp Table 5). However, we found that both of the expressed GADD45A transcripts were bound by multiple ribosomes (Fig 3B). This suggests that while GADD45A is a direct target, many of the transcripts are efficiently translated, similar to what we observed for transcripts with long 3’ UTRs that are NMD targets (Fig 2B).

**Fig 3.**
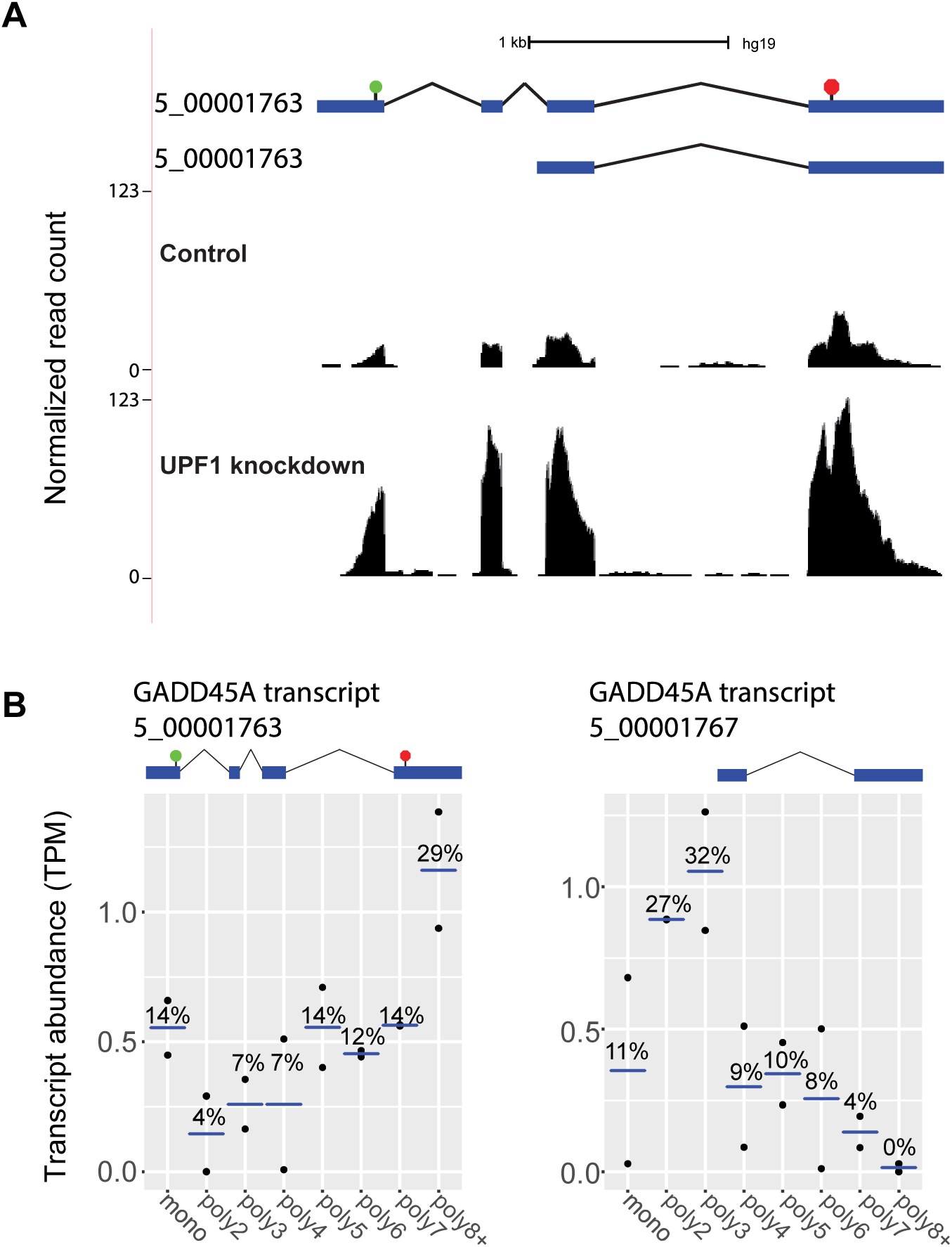
An NMD-targeted transcript responsible for animal lethality is not most abundant in the monosome fraction. (A) The GADD45A transcript isoform 5_00001763 is responsive to NMD inhibition (Supp Table 5). NMD inhibition data was generated with the UCSC Genome Browser. Normalized read counts (# reads/10 million mapped reads) for control cells (top track) and NMD-inhibited cells (bottom track) were plotted. (B) The abundance of the two expressed GADD45A transcripts in the polysome fractions (graphical conventions follow Fig 1). GADD45A (ENSG00000116717) transcript IDs from left to right: 5_00001763 and 5_00001767.

### Transcripts with upstream ORFs and a consensus Kozak sequence often have high abundance in the monosome fraction

Many upstream open reading frames (uORFs) have been identified as features that target a transcript to NMD or translational repression (Wang & Wessler, 1998; Mendell et al., 2004; Yepiskoposyan et al., 2011; Rayson et al., 2012; Son et al., 2017; Ishitsuka et al., 2019), likely due to the presence of exon-exon junctions downstream of the uORF’s stop codon. Because not all uORFs are efficiently translated, it is difficult from sequence analysis alone to identify whether a uORF acts to repress a transcript by altering translation, stability, or both. By examining polysome fractionation data, we can gain insights into whether a uORF-containing transcript is affecting translation, including via the activity of NMD. We examined two well characterized targets of NMD (CHOP and ATF4) that are recognized for degradation through the translation of their uORFs (Mendell et al., 2004; Gardner, 2008; Wang et al., 2011; Karam et al., 2015). In both cases, these transcripts are most abundant in the monosome fraction (Fig 4A), consistent with either these transcripts being translationally repressed, being targeted to NMD, or both. Transcriptome-wide analysis reveals that uORF-containing transcripts are more abundant in the monosome fraction, which is in contrast to transcripts without a uORF (Fig 4B). This is most striking for transcripts whose uORF start codon has a consensus Kozak sequence and is therefore in a strong context for the initiation of translation. Indeed, 12% of transcripts without a uORF are most abundant in the monosome fraction compared to 31 % of transcripts with a uORF in a Kozak sequence (Chi-squared p = 6.29 × 10^−39^; Fig 3B). Transcripts with a uORF start codon not in a Kozak sequence were much less enriched in the monosome fraction with only 15% for with peak abundance in the monosome fraction (Chi-squared p = 6.86 ×10^−20^ when compared to no uORF transcripts; Fig 4B). This is not surprising, given that translation of the uORF is a prerequisite for it to target a transcript to NMD. Taken together, these data suggest that when uORFs act to target a transcript to NMD, they often lead to high abundance in the monosome fraction. Polysome fractionation data could have a broader role in identifying uORFs that exert a regulatory effect on the transcripts via NMD.

**Fig 4.**
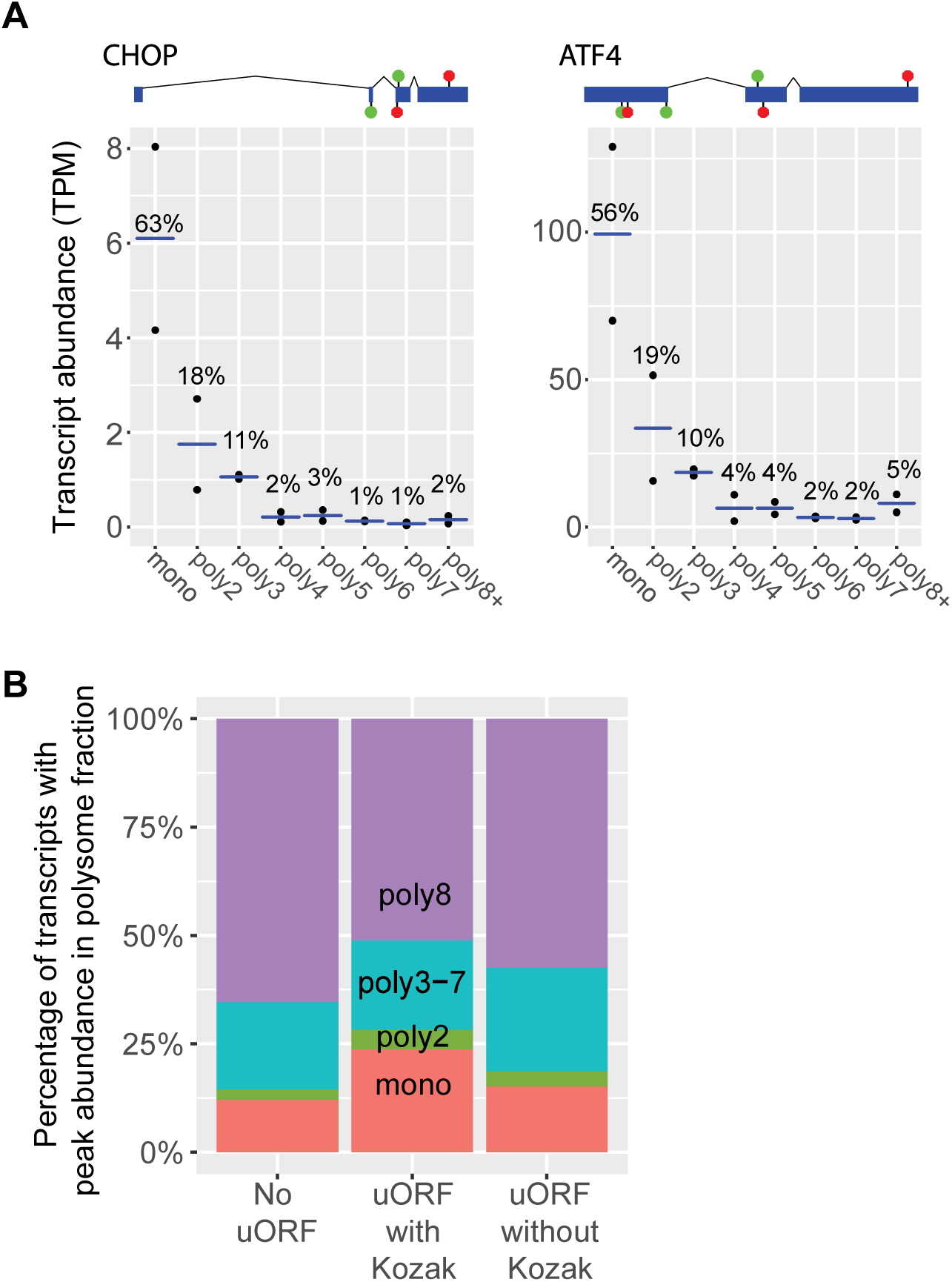
uORF-containing transcripts with a Kozak sequence are enriched in the monosome fraction. (A) Example transcripts whose uORFs have previously been demonstrated to target these transcripts to NMD (graphical conventions follow Fig 1). From left to right, CHOP (ENSG00000175197) transcript ID: 5_00101020 and ATF4 (ENSG00000128272) transcript ID: 5_00152708. (B) Comparison of 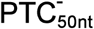 transcripts with and without a uORF according to their peak abundance polysome fraction. uORF-containing transcripts were sub-divided into those with the start codon of the uORF in a Kozak sequence and those without a Kozak sequence.

### Most transcripts annotated as non-coding associated with ribosomes have peak abundance in the monosome fraction and many are likely NMD targets

A growing number of transcripts annotated as non-coding RNAs are being assigned functions; however, the fate of the majority of these transcripts is still unclear (Palazzo & Lee, 2015; Thiel et al., 2019). There are reported examples of non-coding RNAs being targeted to NMD (Kurihara et al., 2009; Smith et al., 2014; Colombo et al., 2017; Li et al., 2017; Zeng et al.,2018), despite the general presumption that RNAs not encoding proteins are not translated. Some annotated non-coding transcripts appear to generate short peptides (Ingolia et al., 2011; Magny et al., 2013; Bazzini et al., 2014; Ingolia et al., 2014; Fesenko et al., 2019). We examined if any transcripts annotated as “non-coding” were found in the polysome fractions, and whether this could be informative regarding their NMD-sensitivity. We compared the proportion of the transcript types across the different polysome fractions. Approximately 15% and 35% of protein-coding 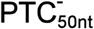 transcripts and 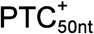 transcripts, respectively, are most abundant in the monosome fraction (Fig 5A). However, for predicted non-coding transcripts, we see a much larger proportion are most abundant in the monosome fraction; 60% for pseudogenes, 52% for long non-coding intergenic RNAs (lincRNAs), 61% for antisense noncoding transcripts, and 57% for other non-coding RNAs (“processed transcripts”) (Fig 5A). The high monosome fraction abundance of pseudogenes is not surprising given that pseudogenes have been reported as NMD targets (Mitrovich & Anderson, 2005; Colombo et al., 2017; Muir et al., 2018). Short ORFs, including those from near-cognate start codons, likely account for the translation of these non-coding transcripts (Guttman et al., 2013). Using previously released analysis of transcript changes upon NMD inhibition in human cells (French, 2016) (Supp Table 5), we found that a majority of non-coding transcripts that are most abundant in the monosome fraction have increased expression upon NMD inhibition (Supp Fig 2).

**Fig 5.**
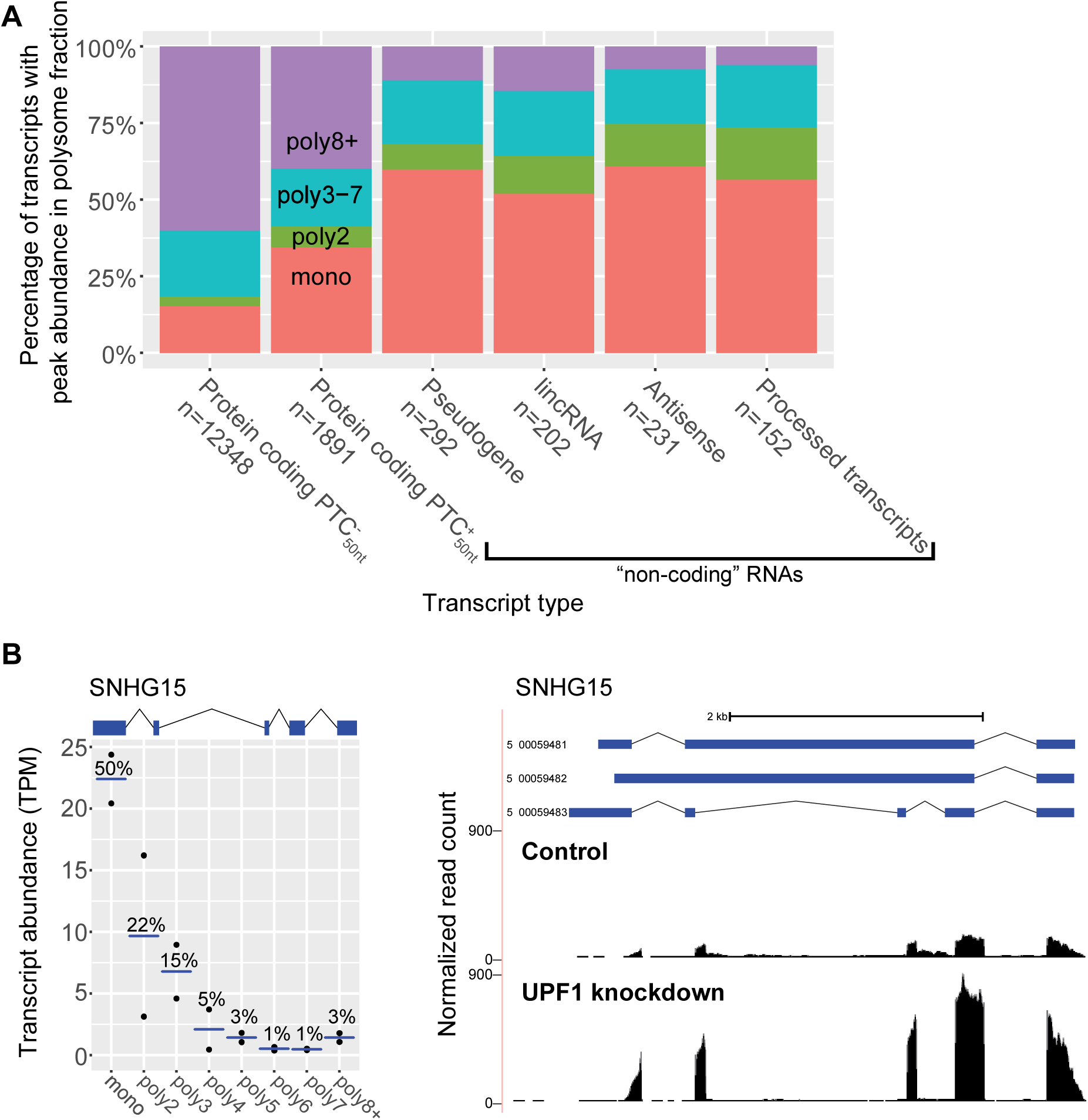
A majority of transcripts annotated as non-coding, that are associated with ribosomes, are most abundant in the monosome fraction. (A) Comparison of transcript types, showing to their peak abundance polysome fraction. Transcript type is defined by the GENCODE gene type, and then further defined by the presence or absence of a PTC_50nt_ within the protein-coding transcripts. (B) Cancer-associated lincRNA SNHG15 (ENSG00000232956; transcript ID: 5_00059483) is an NMD target (graphical conventions follow Fig 1). NMD inhibition data was generated with the UCSC Genome Brower. Normalized read counts (# reads/10 million mapped reads) for control cells (top track) and NMD-inhibited cells (bottom track) were plotted.

We explored in detail SNHG15, which is annotated as a lincRNA and its three transcripts are candidates for being degraded by NMD. SNHG15 expression is associated with poor prognosis in gastric cancer and promoting cancer cell proliferation (Chen et al., 2016; Saeinasab et al., 2019). We find that SNHG15 is most abundant in the monosome fraction and increases in expression when NMD is inhibited (Fig 5B). Previously, the expression of SNHG15 was found to be responsive to the translation inhibitor cycloheximide (Tani & Torimura, 2013, 2015), consistent with SNHG15 being an NMD-targeted transcript. Taken together, these data show that when predicted non-coding transcripts associate with ribosomes, the majority are only bound by a single ribosome and many of these transcripts are NMD-sensitive, including the oncogenic SNHG15.

### Retained intron transcripts are highly abundant in the monosome fraction

We examined if different types of alternative splicing events could affect the abundance of a transcript across the polysome fractions. It has previously been demonstrated that transcripts with retained introns are often localized to the nucleus (Gohring et al., 2014; Boutz et al., 2015; Braun et al., 2017). We found that the splice type with the largest proportion of transcripts in the monosome fraction was intron retention (Fig 6A). As expected, retained intron 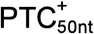 transcripts are highly abundant in the monosome fraction (Fig 6A). To our surprise, 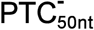 transcripts were also highly abundant in the monosome fraction. Of the 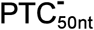 transcripts with retained introns, 22% are most abundant in the monosome fraction (Fig 6A). In contrast, only 11% of 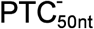 transcripts with cassette exons are most abundant in the monosome fraction (Chi-squared test p = 1.42 × 10^−13^). One possible explanation for the enrichment of retained intron transcripts in the monosome fraction is that many of these transcripts are targeted to NMD, despite the absence of a PTC_50nt_. To test this, we examined the expression change of 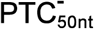 transcripts whose expression is most abundant in the monosome fraction, upon NMD inhibition (French, 2016) (Supp Table 5). We found that 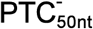 transcripts with cassette exons and those with retained introns have similar responses to the inhibition of NMD (Wilcoxon rank sum test p = 0.81; Fig 5B). Therefore, it is unlikely that NMD is leading to the higher than expected abundance of retained intron transcripts in the monosome fraction, meaning another mechanism is likely involved.

**Fig 6.**
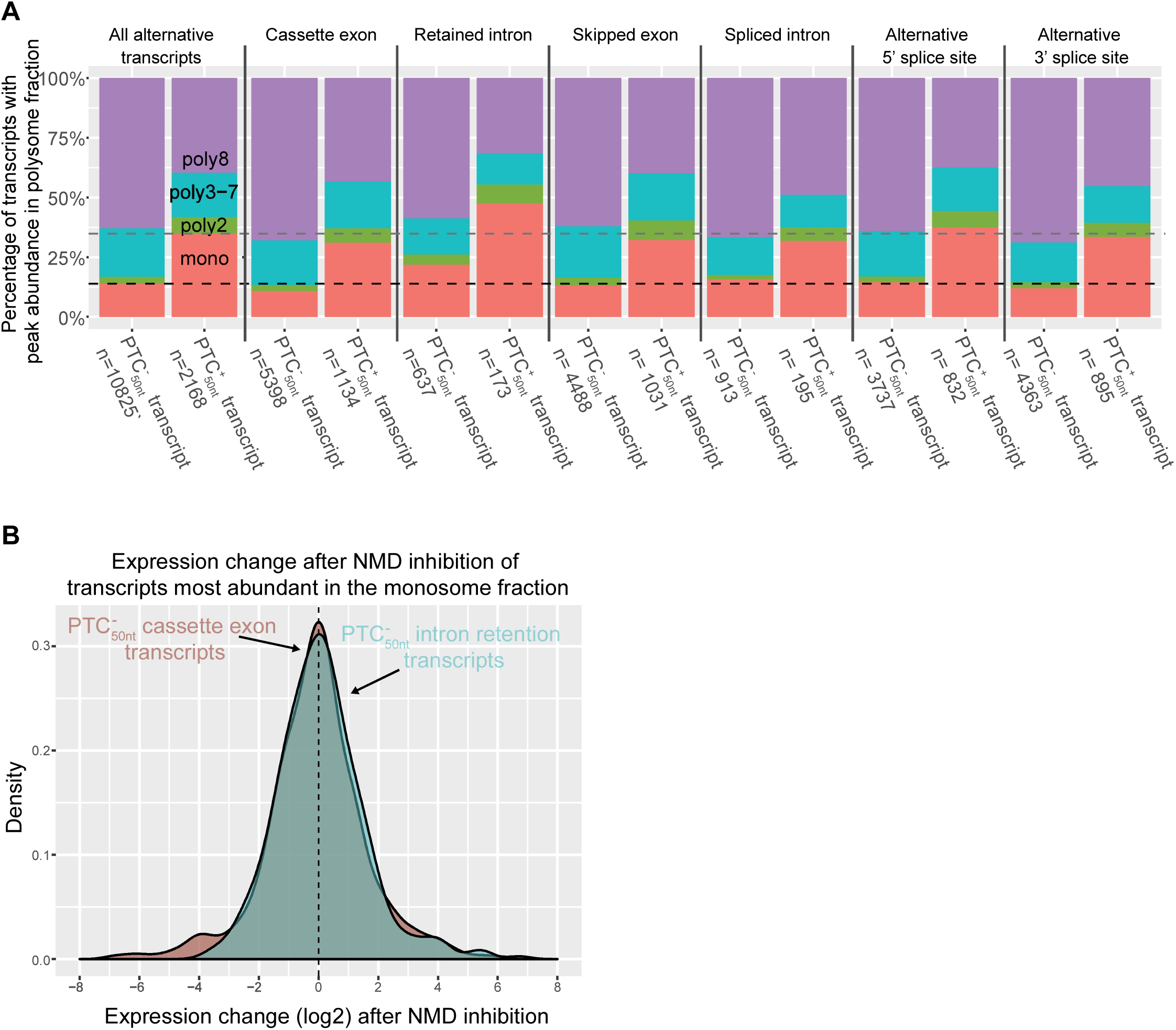
Retained intron transcripts are enriched in the monosome fraction relative to other splice types. (A) Comparison of transcripts with different alternative splicing types according to their peak abundance polysome fraction. Dashed lines represent the average percentage of transcripts with a PTC_50nt_ (grey) and without a PTC_50nt_ (black) that are most abundant in the monosome fraction. (B) The expression change after NMD inhibition of cassette exon or retained intron containing 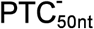 transcripts who are most abundant in the monosome.

## Discussion

NMD is a pervasive mechanism, influencing the expression of thousands of genes in humans (Mendell et al., 2004; Colombo et al., 2017); however, it is not a trivial matter to determine which transcripts are direct NMD targets and what features of a transcript trigger NMD. Studying the steady-state expression of transcripts after NMD inhibition has shown that exon-exon junctions located ≥50 nt downstream of a stop codon are potent triggers of NMD, while it is less clear as to what extent long 3’ UTRs act to trigger NMD (Lindeboom et al., 2016; Colombo et al., 2017; Tian et al., 2017; Lloyd et al., 2018; Hoek et al., 2019). By examining mutations in tumors and the corresponding changes in expression of these genes, it was shown that the EJC model was able to account for the majority of expression changes in mutations leading to early stop codons, and found that the length of the 3’ UTR was not a factor (Lindeboom et al., 2016). In single molecule studies of transcripts, exon-exon junctions were found to be a potent trigger of NMD while the long 3’ UTRs tested were not (Hoek et al., 2019). In this study, we used the abundance of transcripts in the different polysome fractions to explore the features of transcripts linked to NMD. Previous studies have shown that NMD targets are abundant in the monosome fraction compared to polysome fractions (Barberan-Soler et al., 2009; Floor & Doudna, 2016; Heyer & Moore, 2016). This is to be expected given that the first ribosome to terminate at a PTC is likely to induce NMD, which inhibits further translation (Muhlrad & Parker, 1999; Sheth & Parker, 2006; Isken et al., 2008). By analyzing transcript abundance across the polysome fractions, we have been able to explore the features that trigger NMD in physiologically normal cells, without the secondary effects of inhibiting the NMD pathway. Through this approach, we found that the presence of a PTC_50nt_ led to a marked increase in peak abundance in the monosome fraction (Fig 1), in agreement with the EJC model (Nagy & Maquat, 1998; Zhang et al., 1998a; Zhang et al., 1998b; Le Hir et al., 2000a; Le Hir et al., 2000b). NMD-targeted transcripts are often low abundance, meaning that in cells with an active NMD pathway NMD-targeted transcripts do not appear to be expressed. Eighty-four percent of identified NMD-targeted transcript isoforms (French, 2016) (Supp Table 5) were not observed in the polysome fractionization experiment. Despite being unable to detect the majority of the previously NMD-targeted transcripts in the polysome fractions, we were still able to find a strong signal for NMD-triggering features (Fig 1 and 4), and therefore we are likely underestimating the potency of NMD-triggering features such as the EJC model and uORFs.

We did not find that longer 3’ UTRs of 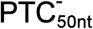 transcripts were associated with higher relative abundance in the monosome fraction (Fig 2). These results suggest that degradation of long 3’ UTR transcripts known to be subject to NMD is not occurring during the pioneer round of translation and therefore these transcripts are not most abundant in the monosome fraction. This would suggest that NMD due to long 3 ‘UTR transcripts has aspects distinct from that of 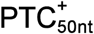 and uORF-containing transcripts.

Approximately half of all human transcripts are computationally identified to contain an uORF (lacono et al., 2005). However, it is difficult to predict whether these uORFs will be translated and if so, whether they will have a repressive effect on the transcript through NMD, translational repression, or both. Consider the two uORF-containing NMD targets, CHOP and ATF4. In normal conditions, these transcripts are targeted to NMD, but when the integrated stress response is activated, both transcripts become highly translated to produce transcription factors integral to the stress response (Gardner, 2008; Wang et al., 2011; Karam et al., 2015; Pakos-Zebrucka et al., 2016). Both transcripts were highly abundant in the monosome fraction (Fig 4A) of this study, consistent with their being targeted to NMD under normal cellular conditions. Globally, we find that a larger fraction of uORF-containing transcripts are most abundant in the monosome fraction than transcripts without a uORF (Fig 4B), consistent with many being translationally repressed or NMD targets or both. Transcripts whose uORF start codon was in a consensus Kozak sequence have larger enrichment in the monosome fraction than do those whose uORF lacks a strong Kozak sequence (Fig 4B).

Many transcripts have been identified in the human genome that appear to lack significant coding potential. These predicted non-coding transcripts come in many flavors, including pseudogenes, lincRNAs and antisense transcripts. Functions have only been assigned to a handful of non-coding transcripts (Palazzo & Lee, 2015; Thiel et al., 2019). Some such transcripts have been identified to produce small peptides (Ingolia et al., 2011; Magny et al., 2013; Bazzini et al., 2014; Ingolia et al., 2014; Fesenko et al., 2019), while others have RNA only functions (Rinn et al., 2007; Gutschner et al., 2011). Using our polysome fractionation analysis, we found that of the predicted non-coding transcripts that do associate with ribosomes, at least 50% in each category of predicted non-coding transcripts are most abundant in the monosome fraction (Fig 5A). Of the non-coding transcripts that are most abundant in the monosome fraction, a majority increase in observed expression when NMD is inhibited (Supp Fig 2), suggesting that these transcripts are direct NMD targets. We discovered that the oncogenic lincRNA SHNG15 appears to be an NMD target, indicating that NMD could be acting to control the expression of physiologically important predicted non-coding transcripts.

Many transcripts with retained introns have been shown to evade NMD because they are not exported from the nucleus (Gohring et al., 2014; Boutz et al., 2015; Braun et al., 2017). We also find ribosome-associated transcripts with retained introns, and these are enriched for greatest abundance in the monosome fraction (Fig 6A, retained intron bars). This would be expected for NMD targets; however, NMD does not appear to explain this large over-representation of retain intron transcripts in the monosome fraction. For instance, retained intron transcripts are also over-represented in the monosome fraction when only intron-retaining 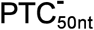 transcripts are considered (Fig 6A). Additionally, the expression of retained intron 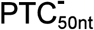 transcripts after NMD inhibition does not indicate that they are NMD sensitive (Fig 6B). Given that uORFs also act as potent triggers for NMD and enrichment in the monosome fraction, we examined 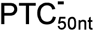 transcripts without uORFs and found that the retained intron were still enriched in the monosome fraction relative to other splice types (Supp Fig 3). It is unclear what is responsible for the high abundance of these transcripts in the monosome fraction, but it is possible other RNA quality control pathways such as No-Go decay or Non-Stop decay (Frischmeyer et al., 2002; van Hoof et al., 2002; Doma & Parker, 2006) could be involved. Alternatively, intronic sequences could act to recruit factors to inhibit translation, activate degradation during translation, or both. Analysis of the translation of splice variants with ribosome profiling found that intron retention events within the coding regions of transcripts were poorly translated (Weatheritt et al., 2016), consistent with our findings that transcripts with retained introns are often only bound by a single ribosome (Fig 6A) and other polysome fractionation studies (Floor & Doudna, 2016; Blair et al., 2017). Further study of the fate of transcript with retained introns will lead to a better understand of the true diversity of roles intron retention plays in gene expression.

In this study, we have shown that using polysome fractionation coupled to RNA-Seq, in a technique called TrIP-Seq (Floor & Doudna, 2016), is a useful tool for the interrogation of the features which trigger NMD in physiologically normal cells. Normally, core NMD factors are depleted, or translation is inhibited, prior to changes in transcript steady state levels being measured to identify NMD-targeted transcripts (Mendell et al., 2004; Drechsel et al., 2013; Hurt et al., 2013; French, 2016; Colombo et al., 2017; Tian et al., 2017; Lloyd et al., 2018). Given that many NMD factors have functions unrelated to NMD (Azzalin & Lingner, 2006; Isken & Maquat, 2008; Lloyd, 2018), indirect changes to the transcriptome could mask the effects of NMD. By measuring transcript abundance in the polysome fractions of physiologically normal cells, without an inhibited NMD pathway, we were able to reveal that transcript features such as a PTC_50nts_ or uORFs, but not long 3’ UTRs, act as potent triggers of NMD. The use of polysome fractionation has the potential to uncover more detailed understanding of the control of gene expression beyond what can be gained from measuring total RNA abundance through RNA-Seq and open up new ways to investigate the fates of transcripts after transcription.

## Materials and methods

### Datasets

We used the polysome fraction RNA-Seq data published by Floor and Doudna (2016) that was deposited in the NCBI SRA database under the GEO Accession ID GSE69352. For NMD inhibition data, we used the data analyzed by French (2016) (https://doi.org/10.6078/D1H019), which can be found in Supp Table 5. The raw reads are deposited in the NCBI SRA database under the BioProject Accession ID PRJNA294972 (SRA Accession ID SRP063462). The transcriptome was a GTF generated by French (2016) (https://doi.org/10.6078/D1H019) that is compatible with the hg19 release of the human genome and can be found in Supp Table 1.

### RNA-Seq analysis

Polysome fraction data from GSE69352 was re-analyzed by mapping to a transcriptome (Supp Table 1) constructed using data from NMD inhibited cells from French (2016) (https://doi.org/10.6078/D1H019). Salmon (Patro et al., 2017) version 0.6.1 was used to quantify these transcripts from the transcriptome FASTA file. The --biasCorrect option was applied and library type was set to -I Ul (inwards paired end reads and unstranded).

Transcript abundance in each sample (measured in Transcript per million; TPM) underwent a bespoke between-sample normalization; the relative amount of RNA in each fraction was estimated from the polysome fractionation experiment, taking into account the proportion of the fraction which was rRNA and non-rRNA molecules. The relative RNA abundance in each polysome fraction was estimated from a polysome profile (Figure 1—figure supplement 1 A of Floor and Doudna, 2016). By measuring the area under each curve of each fraction with ImageJ (Schneider et al., 2012), we were able to estimate the relative amount of total RNA in each fraction (Supp Table 2).

Different polysome fractions will contain different proportions of rRNA and non-ribosomal RNA given the differing numbers of ribosomes bound to a transcript in each polysome fraction. Therefore, we normalized the relative mRNA abundance based on the expected proportions of mRNA in the fraction, as follows. The average length of non-ribosomal RNA molecules was estimated by taking the median length of transcripts (Supp Table 1). This was weighted by the abundance of each transcript, using the raw TPM value from Salmon (Patro et al., 2017). The total length of human rRNA in a ribosome is 7216 nt (sum of GenBank accessions of human rRNA: NR_023379, NR_003285, NR_003287, and NR_003286). We then estimated the proportion of rRNA to non-ribosomal RNA molecules for each fraction with these RNA lengths as follows: For the monosome fraction, it should be one ribosome on each non-ribosomal RNA transcript. However, approximately half of monosomes are empty (Liu & Qian, 2016), therefore we estimate two ribosomes to each transcript. The same ratio is applied to the polysome 2 fraction. The ratio of three to one for the polysome 3 fraction. Four to one for the polysome 4 fraction and so on until the polysome 7 fraction. For the polysome 8+ fraction, we used the estimate used by Floor and Doudna (2016) of, on average, 24 ribosomes to one transcript. This process yielded transcript abundances normalized based on the amount of RNA in each fraction from the polysome gradient.

### Analysis of transcript abundances across the polysome

For each transcript, the fraction containing the highest (peak) abundance was calculated from the average of the two biological replicates, after between-sample normalization (see above). In Fig 1, the proportion of transcripts with peak abundance in each of the polysome fractions was plotted against the distance between the stop codon and the last exon-exon junction. To mitigate effects of noise, the proportion of transcripts with peak abundance in each of the polysome fractions for each position was calculated as an average within a window of distances. If fewer than 200 transcripts had a particular distance between the stop codon and the last exon-exon junction, then the window was expanded by one, in each direction. This was repeated until at least 200 transcripts were captured in the window of distances to be averaged. No more than 350 transcripts were present in any range of window sizes. In Fig 2, the proportion of transcripts with peak abundance in each of the polysome fractions was plotted against the length of the 3’ UTR. The same windowing approach was taken for the 3’ UTR lengths as for the distance between the stop codon and the last exon-exon junction. For plotting, the R (R Development Core Team, 2011) package ggplots2 (Wickham, 2009) or the python module matplotlib (Hunter, 2007) was used.

When analyzing uORFs, the start codons were characterized as being in a Kozak sequence if the sequence matched the pattern RNNATGGV (where R = A or G and V = A or C or G). For gene type analysis, the gene types were drawn from GENCODE version 19 (Harrow et al., 2012); we linked the gene ID from the NMD assembled transcriptome (Supp Table 1), to the Ensembl gene ID via the UCSC gene names (Rosenbloom et al., 2015).

### Annotation of splicing events

We used the annotation script (generateEvents) of the splicing analysis tool SUPPA (Alamancos et al., 2015) to identify alternative splicing events within our transcriptome (Supp Table 1). Taking a GTF file as the input, generateEvents characterizes all of the alternative splicing events in all of a gene’s transcripts and identifies which transcripts include or exclude a particular splicing event, such as a cassette exon. We then annotated each transcript with the splice types associated with it, and whether it was an inclusion or exclusion event for each type.

### Data and code availability

Datasets and code used in this study are available for download at dash.berkeley.edu available at: https://d0i.0rg/l0.6078/D1ZM39.

## Acknowledgments

We thank Stephen Floor for insightful discussions about this research, Anna Desai and Aashish Adhikari for critical comments on the manuscript, and Andrew Sharo for assistance archive the data. This work was supported by NIH Grant R01 GM071655 to SEB, JPBL was supported by the Center for RNA Systems Biology (NIH Grant P50 GM102706 to Jamie Cate), and CEF was supported by NIH T32 GM007232 and T32 HG000047 training grants, and by the Department of Defense (DoD) through the National Defense Science & Engineering Graduate Fellowship (NDSEG) Program.

## Author Contributions

Conceptualization: JPBL and SEB.

Data analysis: JPBL CEF and SEB.

Project administration: JPBL and SEB.

Writing - original draft: JPBL.

Writing - review & editing: JPBL CEF and SEB.

S.E.B reviewed the penultimate version but not the final submitted version due to an accident injury.

## Notes

https://doi.org/10.6078/D1ZM39

