## Supplemental Figures for "Polysome fractionation analysis reveals features important for human nonsense-mediated mRNA decay"

Supp Fig 1

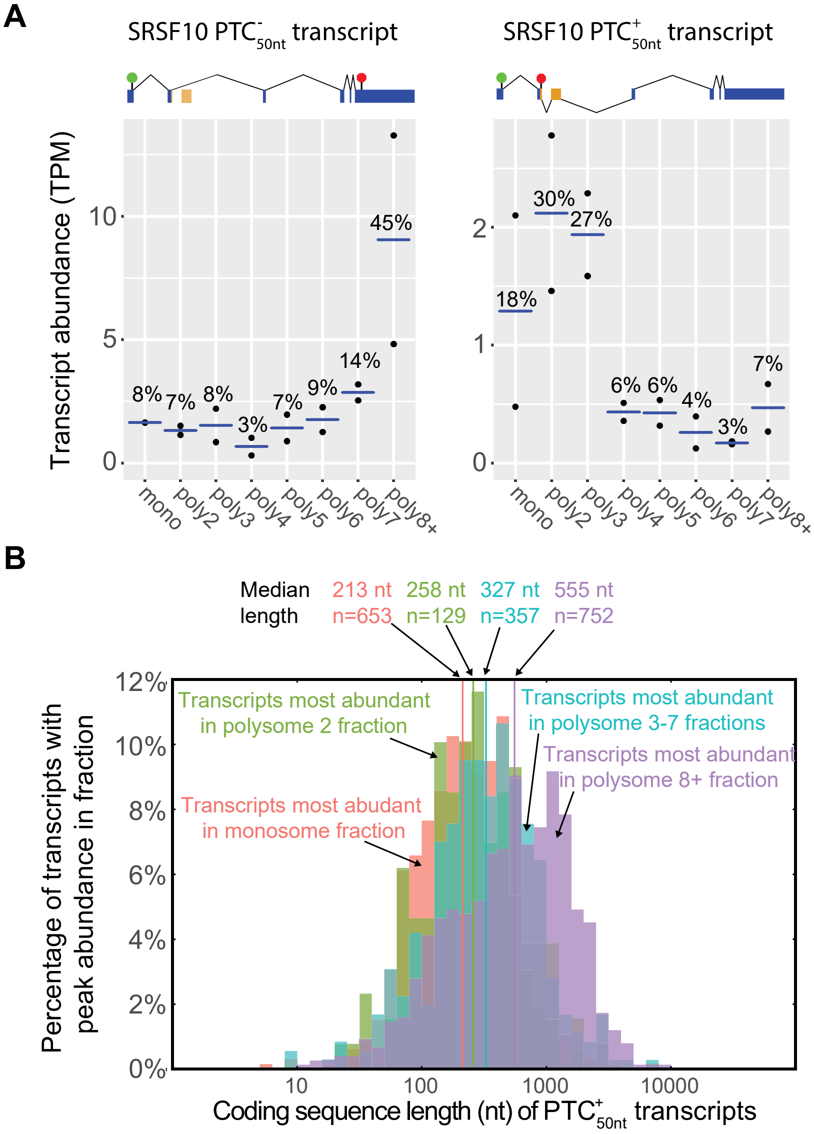
A
SRSF10 PTC$^{-}_{50nt}$ transcript
SRSF10 PTC$^{+}_{50nt}$ transcript
Transcript abundance (TPM)
8% 7% 8% 3% 7% 9% 14% 45%
18% 30% 27% 6% 6% 4% 3% 7%
mono poly2 poly3 poly4 poly5 poly6 poly7 poly8+
B
Median length
213 nt n=653
258 nt n=129
327 nt n=357
555 nt n=752
Transcripts most abundant in polysome 2 fraction
Transcripts most abundant in polysome 3-7 fractions
Transcripts most abundant in polysome 8+ fraction
Transcripts most abudant in monosome fraction
Percentage of transcripts with peak abundance in fraction
Coding sequence length (nt) of PTC$^{+}_{50nt}$ transcripts

Supp Fig 2

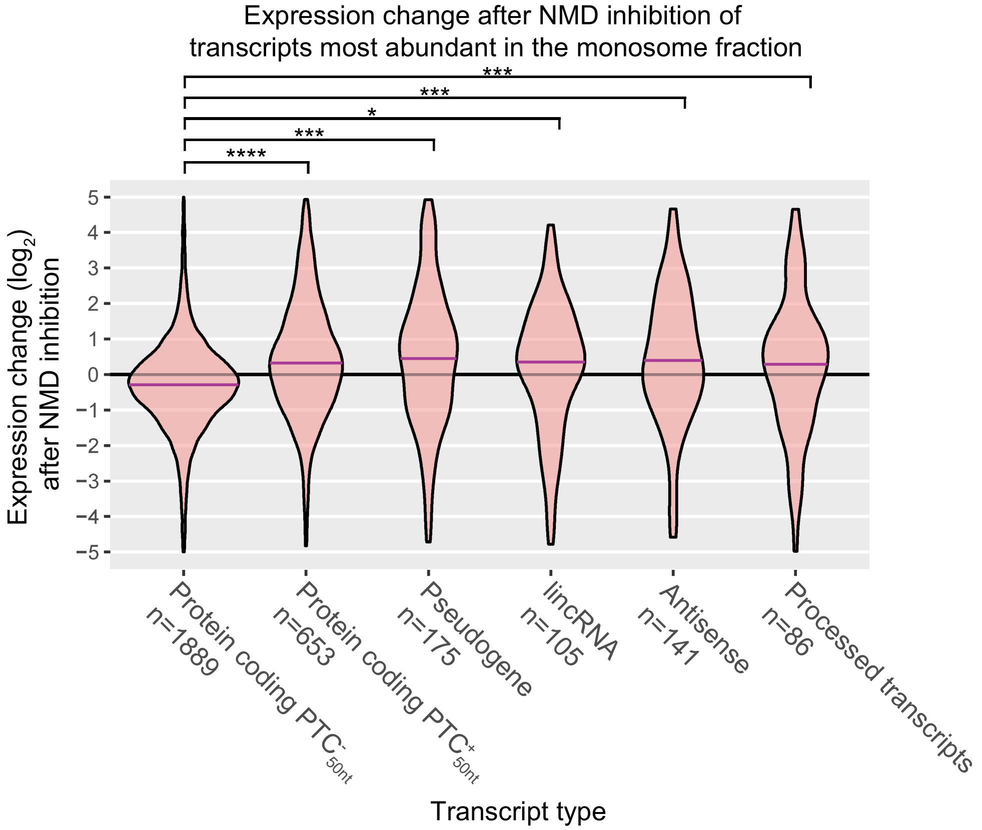
Expression change after NMD inhibition of
transcripts most abundant in the monosome fraction
****
***
*
***
***
5
4
3
2
1
0
−1
−2
−3
−4
−5
Expression change (log2)
after NMD inhibition
Protein coding PTC-50nt
n=1889
Protein coding PTC+50nt
n=653
Pseudogene
n=175
lincRNA
n=105
Antisense
n=141
Processed transcripts
n=86
Transcript type

Supp Fig 3

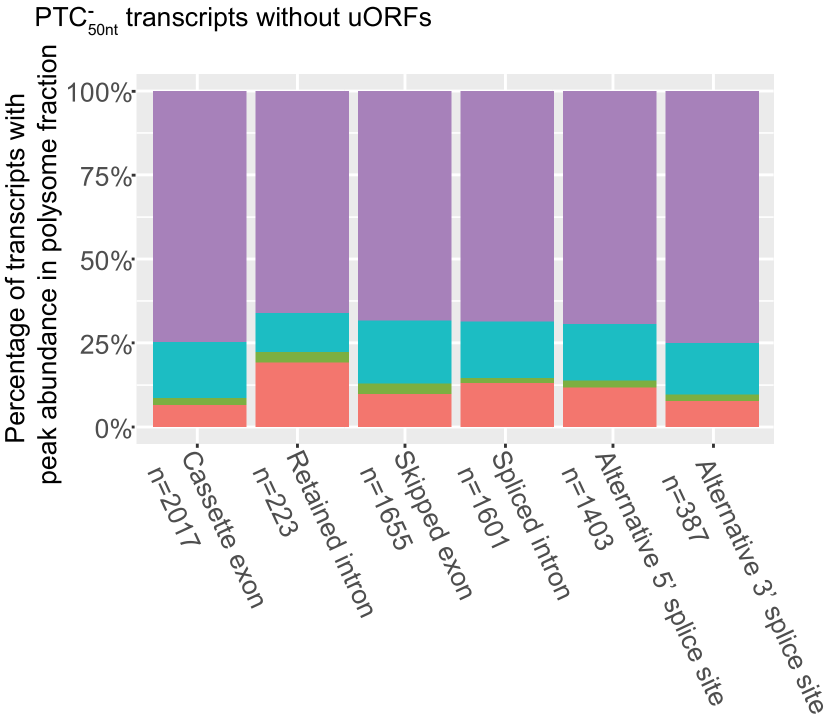
PTC-50nt transcripts without uORFs
Percentage of transcripts with peak abundance in polysome fraction
100%
75%
50%
25%
0%
Cassette exon n=2017
Retained intron n=223
Skipped exon n=1655
Spliced intron n=1601
Alternative 5' splice site n=1403
Alternative 3' splice site n=387

### Supporting information captions

Supp Fig 1. Not all $PTC^{+}_{50nt}$ transcripts are most abundant in the monosome fraction.

(A) SRSF10 transcript isoform abundance profiles. The transcript isoform with a $PTC_{50nt}$ is most abundant in the polysome 2 fraction on average and is also high in the monosome and polysome 3 fractions (graphical conventions follow Fig 1). SRSF10 (ENSG00000188529) transcript IDs from left to right: 5_00005326 ($PTC^{-}_{50nt}$ transcript) and 5_00012473 ($PTC^{+}_{50nt}$ transcript).

(B) The coding sequence lengths of $PTC^{+}_{50nt}$ transcripts versus the peak abundance polysome fraction. On average, the longer the coding sequence, the more ribosomes are observed bound to a $PTC^{+}_{50nt}$ transcripts, although the difference in length of transcripts most abundant in the monosome verse the polysome 2 fractions is not significantly different (average 213nt vs 258nt, Wilcoxon rank sum test p = 0.09).

Supp Fig 2. Expression impact of NMD inhibition on transcripts that are most abundant in the monosome fraction.

The change in expression after NMD inhibition values were taken from (French, 2016) (https://doi.org/10.6078/D1H019) and are presented din Supp Table 5. These values were derived from analysis of transcript expression after UPF1 was knocked down. Transcripts with infinite fold changes upon NMD inhibition were removed from the plot. Wilcoxon rank sum test results: * = p-value <0.01 and ≥0.001, *** = p-value <0.001 and ≥2.2e-16, **** = p-value < 2.2e-16.

Supp Fig 3. $PTC^{-}_{50nt}$ retained intron transcripts are enriched in the monosome fraction relative to $PTC^{-}_{50nt}$ cassette exon transcripts.

Comparison of transcripts with cassette exons or retained introns proportions across the polysome fractions. $PTC^{-}_{50nt}$ transcripts with a uORF have also been excluded from this analysis as this uORF may target the transcript for NMD.

Supp Table 1. Genome annotation file (GTF) of transcript isoforms assembled for hg19 and was used in this study. It was generated from HeLa cells after depletion of UPF1, and is therefore enriched for NMD targets. Reproduced, with permission, from French (2016) (https://doi.org/10.6078/D1H019).

Supp Table 2. Table of individual transcripts and their abundance in the different polysome fractions. This is a tab-delimited text file. Each row represents a different transcript (from Supp Table 1) and columns are identified by the header row: Transcript_ID: Unique transcript code from Supp Table 1, 80S_rep1: Abundance of transcript in the first replicate of the monosome fraction (TPM), 80S_rep2: Abundance of transcript in the second replicate of the monosome fraction (TPM), poly2_rep1: Abundance of transcript in the first replicate of the polysome 2 fraction (TPM), poly2_rep2: Abundance of transcript in the second replicate of the polysome 2 fraction (TPM), poly3_rep1: Abundance of transcript in the first replicate of the polysome 3 fraction (TPM), poly3_rep2: Abundance of transcript in the second replicate of the polysome 2 fraction (TPM), poly4_rep1: Abundance of transcript in the first replicate of the polysome 4 fraction (TPM), poly4_rep2: Abundance of transcript in the second replicate of the polysome 4 fraction (TPM), poly5_rep1: Abundance of transcript in the first replicate of the polysome 5 fraction (TPM), poly5_rep2: Abundance of transcript in the second replicate of the polysome 5 fraction (TPM), poly6_rep1: Abundance of transcript in the first replicate of the polysome 6 fraction (TPM), poly6_rep2: Abundance of transcript in the second replicate of the polysome 6 fraction (TPM), poly7_rep1: Abundance of transcript in the first replicate of the polysome 7 fraction (TPM), poly7_rep2: Abundance of transcript in the second replicate of the polysome 7 fraction (TPM), poly8_rep1: Abundance of transcript in the first replicate of the polysome 8+ fraction (TPM), poly8_rep2: Abundance of transcript in the second replicate of the polysome 8+ fraction (TPM), gene_name: Ensembl gene ID, gene_type: Ensembl gene type, Transcript_length: the length of the transcript (Supp Table 1).

Supp Table 3. Table of sliding window values of stop codon to final exon-exon junction and peak abundance in a polysome fraction. This is a tab-delimited text file. The first column represents the position from the stop codon of the last exon-exon junction of the transcript after the application of a sliding window across each position to ensure at least 200 transcript represent each position, the second column represents the peak abundance of a transcript with that distance of stop to last exon-exon junction. The rows represent transcripts.

Supp Table 4. Table of sliding window values of 3' UTR lengths and peak abundance in a polysome fraction. This is a tab-delimited text file. The first column represents the length of a transcript's 3' UTR after the application of a sliding window across each position to ensure at least 200 transcript represent each position, the second column represents the peak abundance of a transcript with that 3' UTR length. The rows represent transcripts.

Supp Table 5. Table of individual transcripts and their abundance in the different polysome fractions annotated with gene type and splice events present in the transcript isoform. NMD expression values from (French, 2016) (https://doi.org/10.6078/D1H019). This is a tab-delimited text file. Each row represents a different transcript (from Supp Table 1) and columns are identified by the header row: Transcript_ID: Unique transcript code from Supp Table 1, 80S: Percentage abundance of transcript in the monosome fraction, poly2: Percentage abundance of transcript in the polysome 2 fraction, poly3: Percentage abundance of transcript in the polysome

3 fraction, poly4: Percentage abundance of transcript in the polysome 4 fraction, poly5: Percentage abundance of transcript in the polysome 5 fraction, poly6: Percentage abundance of transcript in the polysome 6 fraction, poly7: Percentage abundance of transcript in the polysome 7 fraction, poly8: Percentage abundance of transcript in the polysome 8+ fraction, gene_name: Ensembl gene ID, gene_type: Ensembl gene type, Transcript_length: the length of the transcript (Supp Table 1), Fraction_with_highest_expression: The fraction with the highest abundance of this transcript, POLY_max_expression: Average abundance of the transcript in peak fraction in TPM, NMD_expressed: Whether it is expressed in the NMD inhibition dataset (French, 2016), with an FPKM of at least 1 in one of the two treatments, Strict_NMD_target: Whether a transcript is $PTC^{-}_{50nt}$ (NTC) or $PTC^{+}_{50nt}$ (ETC or PTC), PTC here is a a transcript with a $PTC^{+}_{50nt}$ that was defined by French (2016) as an NMD target, StopCodonToEJ_distance: Distance from stop codon to the last exon-exon junction in nucleotides, AS_types: All the alternative splicing events in this transcript relative to other transcripts of this gene, as defined by SUPPA (Alamancos et al., 2015), AS_inCE: Whether the transcript has a cassette exon included relative to another isoform, AS_inIR: Whether the transcript has a retained intron included relative to another isoform, AS_exCE: Whether the transcript has a cassette exon excluded relative to another isoform, AS_altAA: Whether the transcript has an alternative acceptor site relative to another isoform, AS_altAD: Whether the transcript has an alternative donor site relative to another isoform, AS_exIR: Whether the transcript has a spliced intron relative to another isoform, CF_NMDlog2FC: The change in expression after NMD inhibiotn from French (2016), NTCuORF: Whether the $PTC^{-}_{50nt}$ has a uORF or not, and whether it is in a “strong” or “weak” context as defined by French (2016).
